# Identification of β-synuclein on secretory granules in chromaffin cells and the effects of α- and β-synuclein on BDNF discharge following fusion

**DOI:** 10.1101/158220

**Authors:** Mary A. Bittner, Kevin P. Bohannon, Daniel Axelrod, Ronald W. Holz

## Abstract

Synuclein is strongly implicated in the pathogenesis of Parkinson’s disease as well as in other neurodegenerative diseases. However, its normal function in cells is not understood. The N-termini of α-, β-, and γ-synuclein are comprised of seven 11-amino acid repeats that are predicted to form amphipathic helices. α-Synuclein binds to negatively charged lipids, especially small vesicles and tubulates and vesiculates lipids. The membrane-binding and membrane-curving abilities raise the possibility that synuclein could alter cellular processes that involve highly curved structures. In the present study we examined the localization of endogenous synuclein in bovine chromaffin cells by immunocytochemistry and its possible function to control protein discharge upon fusion of the granule with the plasma membrane by regulating the fusion pore. We found with quantitative immunocytochemistry that endogenous β-synuclein associates with secretory granules. Endogenous α-synuclein only rarely is found on secretory granules. Overexpression of α-synuclein but not β-synuclein quickened the median duration of the post-fusion discharge of BDNF-pHluorin by 30%, consistent with α-synuclein speeding fusion pore expansion.

## Introduction

α-Synuclein was discovered as a protein residing in the presynaptic nerve terminal of the *Torpedo* ray (Maroteaux et al., 1988) and was subsequently demonstrated to be preferentially expressed in mammalian nerve terminals (George et al., 1995; Iwai et al., 1995). α-Synuclein is one of three members of a gene family consisting of α-, β-, and γ-synuclein with 50-60% sequence identity and similar domain organization. Mutations (Kruger et al., 1998; Polymeropoulos et al., 1997; Zarranz et al., 2004) and gene triplication (Singleton et al., 2003) of α-synuclein, but not the other synucleins, are associated with Parkinson’s disease in some patients. A more widespread and direct role for α-synuclein in Parkinson’s disease is suggested by its being a major component of Lewy bodies (Spillantini et al., 1997), which are pathognomonic for Parkinson’s disease and other synucleinopathies. α-Synuclein is also a component of senile plaques in Alzheimer’s disease (Ueda et al., 1993). Although there is strong evidence for a pathogenic effect of synuclein, its role in causing disease and its normal function in cells is incompletely understood [see review (Bendor et al., 2013)].

The presynaptic localization of synuclein raises the possibility that it normally regulates synaptic transmission and secretion. Indeed, α-Synuclein is a modest negative regulator of catecholamine secretion in chromaffin cells (Larsen et al., 2006) and in nigrostriatal (Abeliovich et al., 2000; Yavich et al., 2004), and hippocampal (Nemani et al., 2010) nerve terminals by reducing the frequency of exocytotic events. The mechanism for these effects, their physiological significance and their relationship to synuclein-induced disease is uncertain Synuclein may modulate SNARE interactions that are necessary for exocytosis. It has been reported that α-synuclein promotes SNARE complex formation (Burre et al., 2010) and, in addition, maintains the nerve terminal because of protein chaperone activity (Chandra et al., 2005). How these activities are related to the inhibitory effects of synuclein on exocytosis is unclear. The biochemistry of synuclein raises the possibility of an additional role of synuclein in secretion.

The N-termini of α-, β-, and γ-synuclein are comprised of seven 11-amino acid repeats that are predicted to form amphipathic helices. Indeed, α-synuclein binds to negatively charged lipids, especially small vesicles (Davidson et al., 1998) and tubulates and vesiculates lipids (Westphal and Chandra, 2013). Upon lipid binding, the secondary structure of α-synuclein changes from random to strongly alpha-helical (Davidson et al., 1998). The membrane-binding and membrane-curving abilities raise the possibility that synuclein affects the highly curved fusion pore that arises upon fusion of the granule membrane with the plasma membrane. The notion of a protein acting to regulate fusion pore expansion because of its membrane-bending activity has a strong precedent in our work concerning the membrane-curving protein dynamin, the master fission controller in endocytosis (Anantharam et al., 2011; Anantharam et al., 2010). Another membrane-curving protein, synaptotagmin, also regulates fusion pore expansion (Rao et al., 2014).

In the present study we investigated with several anti-synuclein antibodies the localization α-and β-synuclein in bovine chromaffin cells and the effects of α-and β-synuclein overexpression on protein discharge upon secretory granule fusion. Immunocytochemistry indicates that endogenous β-synuclein, but not α-synuclein, is associated with most chromaffin granules. Overexpressed α-synuclein, but not β-synuclein, quickened the discharge of co-transfected BDNF-pHluorin, consistent with α-synuclein speeding fusion pore expansion. A closely related study was published while this work was in progress (Logan et al., 2017).

## METHODS

### Culture and transfection

Primary bovine adrenal medullary chromaffin cells were isolated as previously described (Wick, 1993), and plated on 35 mm glass-bottom dishes (refractive index = 1.51, World Precision Instruments, Sarasota, FL) treated with poly-D-lysine and layered with bovine collagen. Chromaffin cells were transfected with the Neon Transfection System (Invitrogen). 1 x 10^6^ cells and up to 10 µg DNA in 100 µl total resuspension buffer were electroporated with pulse settings of 1100 mV and 40 ms. BDNF-pHluorin was constructed from BDNF (prBDNF-stop-EGFP-N1), which was obtained from Professor V. Lessman (Otto-Von-Guericke Universitat, Magelburg, Germany). EGFP-labeled human α-synuclein was obtained from Addgene. Unlabeled α-synuclein was constructed by removing the synuclein sequence and inserting it into pcDNA3 using HindIII and EcoRI restriction sites. The construct used to express human β-synuclein was made using synthetic gBlocks (Integrated DNA Technologies, Coralville, IA). The β-synuclein consensus sequence used was from accession number NM_001001502.2. The coding sequence was flanked by HindIII and the Kozak sequence GCCGCCACC on the 5’ end and a stop codon and BamHI on the 3’ end. The gBlock was digested with HindIII and BamHI and ligated into pcDNA3.1(+).

### Immunocytochemistry - Confocal Microscopy

Images were acquired on an Olympus Fluoview 500 confocal microscope with a 60 x 1.42 numerical aperture oil objective. An argon 488 nm laser with a 505–525 nm bandpass filter, a HeNe green 543 nm laser with a 560-600 nm bandpass filter, and a HeNe red (633 nm) laser with a longpass filter were used. To minimize spillover, images with different excitations were acquired sequentially. Within an experiment, initial settings were adjusted so that the brightest pixels for each color were unsaturated, and these settings were maintained throughout. Images were analyzed with ImageJ software (NIH), and statistics were performed with Prism 6 software from Graphpad Prism Software.

### Antibodies

Antibodies were from the following sources: DSHB Hybridoma Product H3C against α/β-synuclein (Developmental Studies Hybridoma Bank, The University of Iowa, Iowa City, IA 52242); antibodies specific for **α**-or **β**-synuclein (Abcam 138501 and Abcam 15532, respectively); Alexafluor™-labeled secondary antibodies (Life Technologies (Molecular Probes)).

### Secretion experiments

All experiments were performed 3-5 days post transfection in a room heated to 34 ± 1°C. Individual cells were perfused through a pipet (100 μm inner diameter) using positive pressure from a computer-controlled perfusion system DAD-6VM (ALA Scientific Instruments, Westbury, NY). The bath solution and the initial perfusion solution were physiological saline solution (PSS, 145 mM NaCl, 5.6 mM KCl, 2.2 mM CaCl2, 0.5 mM MgCl2, 5.6 mM glucose, and 15 mM HEPES, pH 7.4). Other solutions used during perfusion were elevated potassium PSS (KPSS, 95 mM NaCl, 56 mM KCl, 2.2 mM CaCl2, 0.5 mM MgCl2, 5.6 mM glucose, and 15 mM HEPES, pH 7.4) and low pH MES buffer (MES, 145 mM NaCl, 5.6 mM KCl, 2.2 mM CaCl2, 0.5 mM MgCl2, 5.6 mM glucose, and 15 mM MES, pH 5.5). Cells were generally perfused according to the following schedule: 3 s PSS, 3 s MES, 3 s PSS, 45 s KPSS, 5 s MES, 10 s PSS.

### TIRFM

A 488 nm laser (Coherent OBIS or Melles Griot 543-AP-01) was used to excite pHluorin The filter cube contained a dichroic mirror/emission filter combination: ZT488/561rpc and ZET488/561m (Chroma Technology, Brattleboro, VT). Objective-based TIRF illumination was produced by directing the excitation beam to computer-controlled galvanometer mirrors that directed the beam through a custom side port to a side-facing filter cube below the objective turret of an Olympus IX70 (inverted) microscope (Melville, NY). The aligned excitation beams were focused and positioned near the periphery of the back focal plane of a 60x 1.49 NA, oil immersion objective (Olympus) so that the laser beam was incident on the coverslip at ~70° from the normal giving a decay constant for the evanescent field of approximately 100 nm. The galvanometer mirrors were computer controlled through a DAQ board, NI PCIE-6351 (National Instruments, Austin, TX) and a custom LABVIEW program. The cells were imaged at 36 Hz.

### Analysis of event duration

PHluorin-labeled protein discharge was measured over time for a small region of interest (~0.7 μm diameter) centered on the event using Time Series Analyzer V2.0 plugin in Image J. Time-varying local backgrounds were determined by capturing the intensity of a neighboring ROI without a fusion event. They were subtracted frame by frame from the intensities of the discharge events. The local background subtraction was necessary because of increases in background intensity due to protein diffusion from nearby events. The duration of discharge of pHluorin-labeled proteins was determined with a custom program written in IDL that largely eliminated subjectivity of the analysis and greatly facilitated the interpretation of the results, especially with complex intensity vs time profiles. In this program, the user defines the start and end times in the fluorescence vs time curve for each event. The value of the fluorescence at the chosen start time *t*start, just before fluorescence begins to rise, is considered to be the "baseline". The end time *t*end is chosen to be where the fluorescence after the event has returned to its lowest value. The program then determines the time of the maximum fluorescence *t*max within this time window. The intervals (*t*start, *t*max) and (*t*max, *t*end) are defined as the "rise phase" and "fall phase", respectively. Each of those phases is first best-fit with a 5th-degree polynomial (for smoothing), and then with a sloping straight line, upward for the rising phase and downward for the falling phase. Straight lines with those slopes are then "pinned" to the positions of the maximum (upward or downward) slopes of the fluorescence data, and extrapolated to the baseline. The time period between the baseline intercept of the rising phase straight line and the falling phase straight line is considered to be the duration (i.e., characteristic time) of the event.

## RESULTS

### Immunocytochemical detection of α-and β-synuclein in bovine chromaffin cells

We examined the localization of α-and β-synuclein in cultured bovine chromaffin cells by immunocytochemistry using a battery of three antibodies, one specific for α-synuclein, the second specific for β-synuclein, and a third which is non-isoform specific and detects both α-and β-isoforms. The sequences of bovine α-and β-synuclein and the epitopes seen by the antibodies are shown in **Fig. 1**. The specificity of each antibody was verified by its ability to recognize the appropriate overexpressed protein(s).

**Figure 1.**
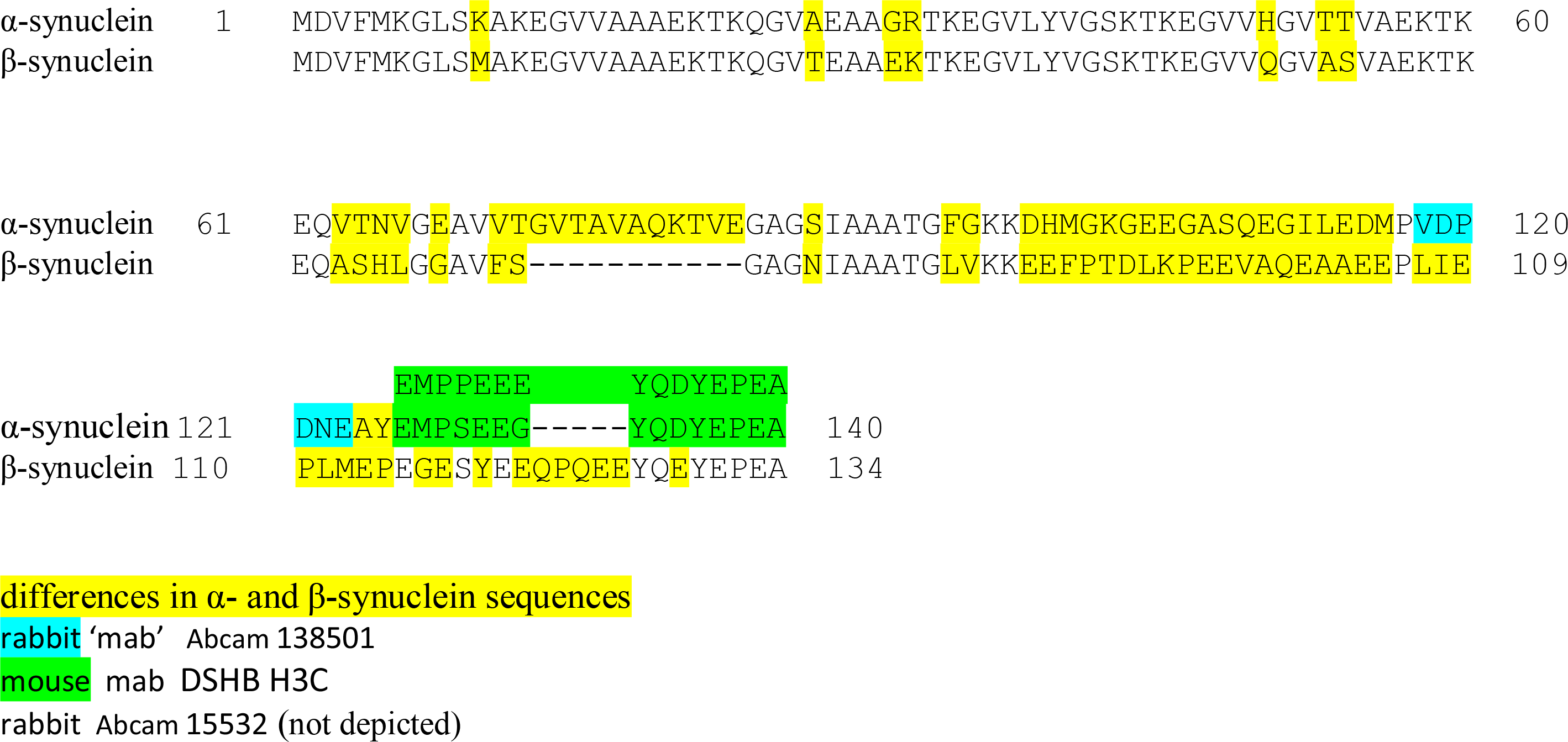
Anti-Synuclein Antibodies and Bovine α- and β-Synuclein Sequences. The rabbit “monoclonal” antibody (Abcam 138501) recognizes only **α**-synuclein (the corresponding sequence in β-synuclein is completely different). The antibody was made against a peptide (aa 118-123 VDPDNE, in blue) in human α-synuclein (identical to bovine sequence). The rabbit antibody (Abcam 15532) recognizes **β**-synuclein only (epitope near C-terminus, not depicted). The mouse monoclonal antibody (DSHB H3C) recognizes both **α**-synuclein and **β**-synuclein. It probably sees the very C-terminal amino acids. The antibody was generated against a peptide (aa 126-140 EMPPEEEYQDYEPEA in green) in canary α-synuclein (shown above bovine sequences). Differences in bovine α- and β-synuclein sequences are highlighted in yellow.

In bovine chromaffin cells, all three anti-synuclein antibodies exhibited punctate rather than diffuse cytoplasmic staining (**Fig. 2**). Puncta labeled by the antibody specific for β-synuclein (**Fig. 2D**) and the antibody that recognizes both α-and β-synuclein (**Fig. 2B**) were abundant, while the puncta labeled by the α-synuclein antibody were relatively sparse (**Fig. 2E**).

When we labeled newly synthesized chromaffin granules by expressing BDNF-pHl, the non-isoform-specific antibody to the common C-terminus of α-and β-synuclein was strongly co-localized with BDNF-pHL, labeling 85 ± 2% of BDNF-pHl granules (**Fig. 2 A & B**, yellow arrowheads). It was also present on numerous other puncta, perhaps granules made prior to transduction with BDNF-pHL. The antibody specific for β-synuclein also recognized a majority of BDNF-pHl granules, labeling 69 ± 1% of them (**Fig. 2 C & D**). In contrast, puncta seen by the α-synuclein antibody rarely coincided with BDNF-pHl granules, labeling only 11 ± 2%. The data indicate that endogenous synuclein is associated with chromaffin granules, with β-synuclein the major isoform in bovine cells.

**Figure 2.**
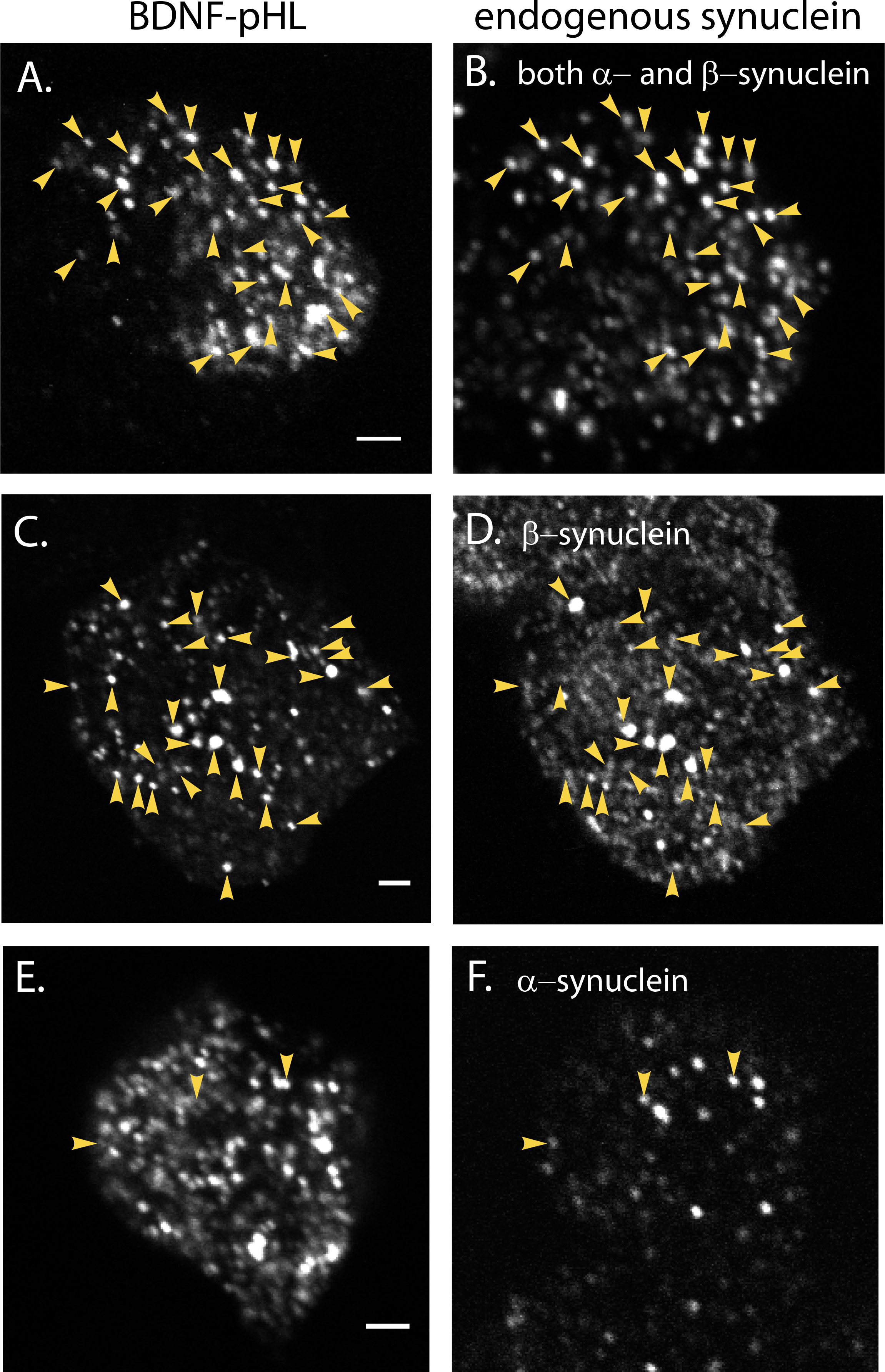
Co-localization of endogenous synuclein with chromaffin granules. Bovine chromaffin cells transiently expressing the chromaffin granule marker BDNF-pHL were fixed with 4% paraformaldehyde, permeabilized with methanol, and incubated with various antibodies that recognize either or both of the α- and β-isoforms of bovine synuclein. Cells were then rinsed and incubated with a 2° antibody (Alexafluor647™-labeled donkey anti-mouse IgG). Cells were imaged on a confocal microscope (60x magnification) as described in *Methods*. (A, C, E) Newly synthesized chromaffin granules containing BDNF-pHL; (B) endogenous synuclein as detected by DSHB H3C, an antibody which recognizes both α- and β-synuclein. Arrowheads indicate some of the co-localized puncta. Synuclein puncta in a neighboring non-transfected cell are visible at the bottom of the image. (D) endogenous synuclein as detected by an antibody specific for β-synuclein (Abcam 15532); (F) endogenous synuclein as detected by an antibody specific for α-synuclein (Abcam 138501). Scale bars = 2 µm.

### Effects of overexpressed of α-or β-synuclein on discharge of BDNF-pHluorin from chromaffin granules

Chromaffin cells were co-transfected with a plasmid encoding BDNF labeled with the highly pH sensitive GFP, pHluorin (pHl) and either a plasmid encoding α-or β-synuclein or control plasmid (pcDNA3). Immunocytochemistry indicated that transfected α-synuclein or β-synuclein was co-expressed with BDNF-pHl in 95-100% of the cells.

PHluorin is almost undetectable in the acidic interior (pH ~5.5) of the secretory granule in living cells before fusion. Upon fusion there is a profound increase in pHluorin fluorescence as the granule lumen equilibrates with the extracellular pH (7.4) followed by a decrease in intensity as the protein diffuses into the medium (**Fig. 3A**). Both the rate of the initial increase in fluorescence and the time course of its subsequent decrease as the protein diffuses away provide indirect information about fusion pore expansion. The kinetics of labeled protein discharge proved to be complex, with variable shapes of fluorescence intensity vs time and uncertain baseline plateaus. Therefore, software to quantitatively describe the duration of the events should rely, not on absolute intensity above baseline, but rather on rising and falling slopes of intensity. Our method, written in a custom IDL program, is robust against changes in event shape and is relatively insensitive to baseline selection. **Fig. 3A** shows a relatively simple intensity profile. The increase in fluorescence occurred within two frames (56 ms). The falling phase was initially very rapid followed by a long plateau. The program, by taking an average of the falling slope (weighted by the relative intensity) calculated a duration of 4.6 sec (see method for details).

**Figure 3.**
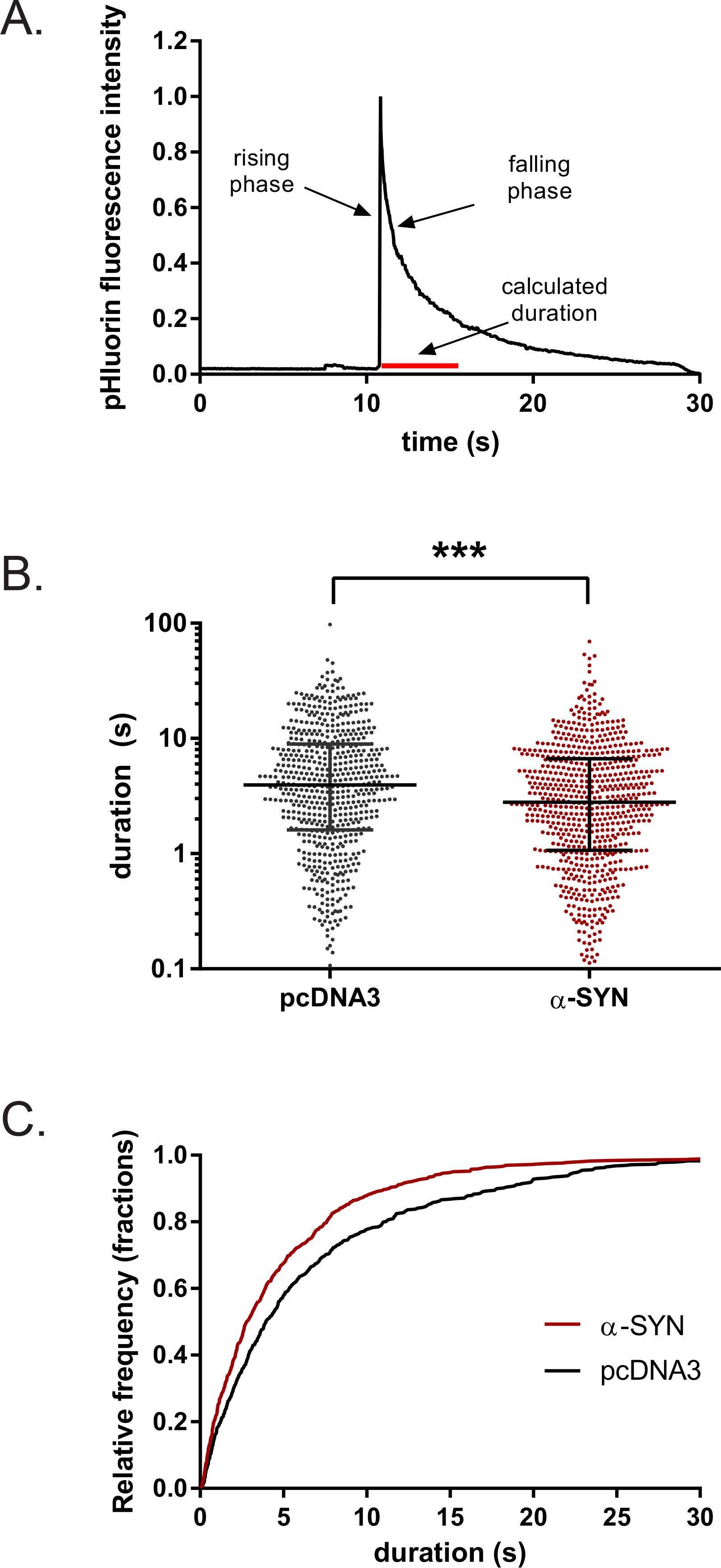
αSynuclein speeds the release of BDNF-pHl from bovine adrenal chromaffin cells. (A) An example of the discharge of BDNF-pHl from a secretory granule upon fusion. The initial increase in fluorescence was too rapid to resolve with imaging at 36 Hz (less than 56 ms). The estimated duration, 4.6 s (red bar), was determined by custom software (see text). (B) Scatter plot of event durations. Horizontal bars indicate the median (longest bar) and the upper and lower quartiles. Note that the ordinate is in log scale. ***, p < 0.0001 as determined by a Kolmogorov-Smirnov test. (C) Cumulative histogram of BDNF-pHl event durations in the absence and presence of α-synuclein from panel B.

Live cell experiments were conducted at 34-35°C, close to physiological temperature with imaging at 36 Hz. Cells were stimulated by perfusion of single cells with solution containing elevated K^+^ (see Methods for details). α-Synuclein quickened the post-fusion discharge of BDNF-pH. Median duration decreased from 3.9 (593 events) to 2.8 s (685 events) (**Fig. 3B**). The cumulative distributions were significantly different (Kolmogorov-Smirnov test, p < 0.0001) (**Fig. 3C**). In contrast, β-synuclein did not have a significant effect on BDNF-pHl discharge (**Fig. 4**). The median duration of BDNF-pHl in the presence of β-synuclein (101 events) was 3.6 s, compared to 3.0 s for the pcDNA3 control group (41 events).

**Figure 4.**
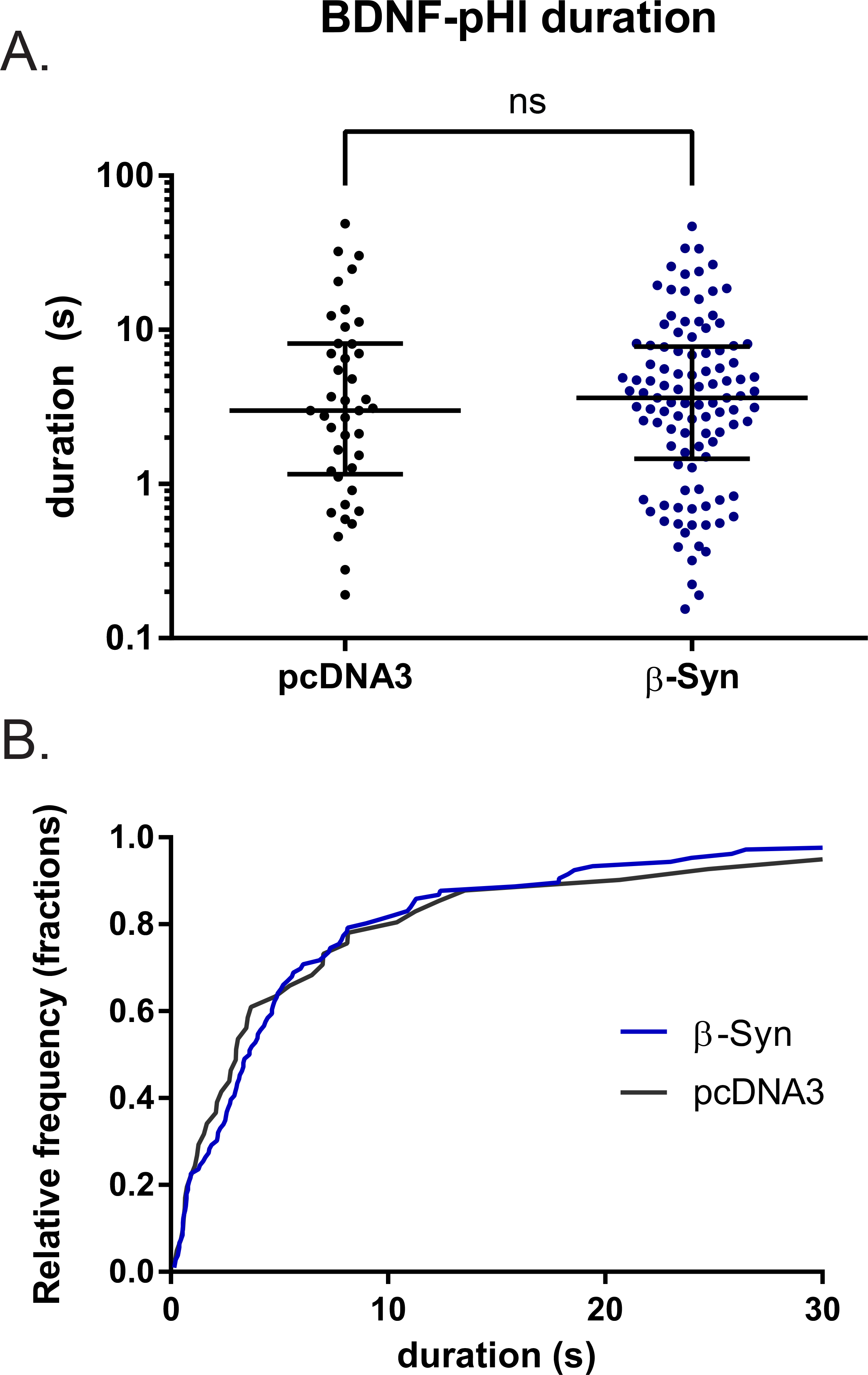
βSynuclein does not speed the release of BDNF-pHl from bovine adrenal chromaffin cells. (A) Scatter plot of event durations. Horizontal bars indicate the median and the upper and lower quartiles. Note that the ordinate is in log scale. ns, p = 0.91 as determined by a Kolmogorov-Smirnov test, (B) Cumulative histogram of BDNF-pHl event durations in the absence and presence of β-synuclein from panel A.

Most BDNF-pHl events reached peak fluorescence within one or two frames (28-56 ms). We could not detect a change in the rise times upon overexpression of either α-or β-synuclein.

## DISCUSSION

There are two main conclusions from our work. First, β-synuclein but not α-synuclein is the synuclein isoform most associated with secretory granules (chromaffin granules) in chromaffin cells. Both an antibody specific for β-synuclein (Abcam 15532) and one that detects both α-and β-synuclein (mouse mab DSHB H3C) detected antigen on 70-85% of newly synthesized secretory granules (labeled with transfected BDNF-pHluorin). An antibody specific for α-synuclein (Abcam 13850) only rarely detected antigen on these granules. The same α-and β-synuclein-detecting antibody DSHB H3C also recognizes secretory granules in mouse chromaffin cells (Logan et al., 2017). The synuclein isoform was not identified in the mouse study.

We also examined the antibody that Tompkins *et al* (Tompkins et al., 2003) used (Syn1, S63320) in bovine cells, and we confirm their results. The antibody did not recognize bovine α-synuclein on granules. It did recognize an antigen in Golgi as the paper indicated. In a subsequent investigation (Perrin et al., 2003), the epitope was mapped to residues 91-99 in human α-synuclein. The bovine sequence (91-99) has 2 amino acid differences that could render the antibody unable to recognize bovine synuclein. It was found that the antibody also recognizes a 45 kDa protein that is present in α-synuclein null mice. We suspect that the Golgi antigen seen in bovine chromaffin cells is this 45 kDa protein.

Second, overexpression of α-synuclein, but not β-synuclein decreased the discharge time for transfected BDNF-pHl following fusion. The effect was relatively modest with a reduction of the median discharge time of 30%, from 3.9 s to 2.8 s. The speed of protein discharge after fusion is assumed to be a surrogate for fusion pore expansion and may therefore reflect an effect of α-synuclein to cause more rapid fusion pore expansion. These results are in agreement with those of Logan et al. (Logan et al., 2017) in which α-but not β-synuclein increased post-fusion discharge rates of BDNF-pHl. Since endogenous β-but not α-synuclein is localized to chromaffin granules, the physiological significance of the α-synuclein effect is uncertain. Protein discharge was significantly slowed in chromaffin cells derived from mice without α-, β-, γ-synuclein (triple knockout)(Logan et al., 2017) consistent with a role of one or more of the endogenous synucleins in regulating post-fusion protein discharge. Experiments were not performed to determine whether the wild type phenotype could be rescued by infection with virus expressing a synuclein isoform.

The rise time of the BDNF-pHl fluorescence upon fusion reflects the initial time course of fusion pore expansion convoluted with neutralization of the highly buffered lumenal interior. The rise times with and without transfected α-synuclein were faster than the time resolution of our experiments (~56 ms). In contrast, Logan et al (with a lower time resolution, ~100 ms) detected rise time of 200-300 ms. α-Synuclein quickened rise times. The more physiological temperature in our experiments (34-35°C) compared to room temperature in the experiments of Logan et al. probably resulted in rise times too rapid to be resolved.

## Acknowledgements

We thank Annita Ngatchou Weiss for initiating the synuclein experiments in the laboratory and Prabhodh Abbineni for many helpful discussions. We are grateful to William T. Dauer for providing synuclein constructs and timely advice. We thank Robert H. Edwards and Todd Logan (UCSF) for engaging discussions and sharing their prepublication findings. Our work was supported by Rapid Response Innovation Award from the Michael J. Fox Foundation for Parkinson’s Research to RWH, by NIH Grant RO1-170553 to RWH and DA, and by 5T32HL7853-17 to KPB. The hybridoma H3C developed by J. George was obtained from the Developmental Studies Hybridoma Bank, created by the NICHD of the NIH and maintained at The University of Iowa, Department of Biology, Iowa City, IA 52242.

